# CyranoPS, a novel compact temperate corynephage infecting *Corynebacterium glutamicum* MB001

**DOI:** 10.64898/2026.10.29.762952

**Authors:** Biel Badia Roigé, Samy Mechrouki, Elsa Chaouat, Océane Hill, Nathan Beaure, Julia Frunzke, Christophe Regeard, Ombeline Rossier

## Abstract

Here, we describe CyranoPS, a temperate phage isolated from the river Yvette in France, using the prophage-free strain *Corynebacterium glutamicum* MB001. Negative-stain electron microscopy revealed virions with siphovirus morphology. Its 14,910-bp genome has a GC content of 53.0% and 14-nt 3’ overhangs at both ends. It carries 26 predicted genes, 10 without predicted functions, and no tRNAs. CyranoPS belongs to the newly established actinophage cluster ER, whose members have the smallest known genomes among *Corynebacterium*-infecting phages. Deletion of genes encoding the porin PorAH and the protein mycoloyltransferase MytC abolished infection, while genes encoding the mycomembrane proteins PorBC and ProtX were dispensable. CyranoPS established lysogeny in *C. glutamicum* through chromosomal integration probably mediated by its predicted tyrosine integrase. Integration truncated the immunity repressor gene Cyr_18, resulting in a protein devoid of its 15 C-terminal residues. The truncated repressor conferred stronger protection against superinfection than the full-length repressor. CyranoPS lysogens exhibited high levels of spontaneous prophage induction, reaching 10^9^ PFU/mL under standard conditions, with no further increase upon treatment with mitomycin C. These findings suggest a potentially SOS-independent lytic-lysogenic switch and establish CyranoPS as a compact model to study infection and lysogeny regulation in Mycobacteriales.

## Introduction

Bacteriophages, or phages for short, are bacterial predators found throughout every environment on Earth. As the most abundant biological entities on the planet, an estimated 10^31^ phage particles exist in nature, driving an estimated 10^24^ infection events per second (Clokie et al. 2011; Cobian Guemes et al. 2016). These events shape bacterial community structure and fuel an ongoing evolutionary arms race between phages and their hosts (Hendrix et al. 1999; Hendrix 2002).

While phages offer valuable tools for biotechnology, they also pose a threat to biotechnological production processes: phage contamination in industrial fermentations can cause biomass loss and, consequently, economic and production losses (Kaminski and Paczesny 2024). *Corynebacterium glutamicum* is a key biotechnological model organism; the strain *C. glutamicum* ATCC 13032 (NCBI accession number BX927147) in particular is used industrially for amino acid production (notably more that 5 million tons of glutamate and lysine per year) and serves as a platform strain for producing a range of other compounds (Becker et al. 2018; Kalinowski et al. 2003). Its genome carries two small, degenerated cryptic prophages, CGP1 and CGP2, and a defective integrative and conjugative element, CGP3 (ca. 187 kb), which can excise from the genome upon SOS response induction (Badia Roige et al. 2026; Frunzke et al. 2008a; Pfeifer et al. 2016). Although most of the CGP3 region remains uncharacterized, it is known to encode several anti-phage (or phage-interacting) elements, including a restriction modification system (Schafer et al. 1997) and the xenogeneic silencer CgpS (Pfeifer et al. 2016).

Phages infecting corynebacterial species are known as corynephages. While the first phages infecting *C. glutamicum* strains were described as early as the 1970s, the set of phages known to infect *C. glutamicum* ATCC 13032 specifically remains limited. Recent work on phages CL31 and Cog has highlighted the importance of the mycolic acid layer for adsorption and successful infection and shown that the RM system encoded in CGP3 is critical for restricting CL31 infection (Hunnefeld et al. 2021; McKitterick and Bernhardt 2022). Recently described phages infecting this strain are PSonyx (Rossier et al. 2024), MicyPS (Rossier et al. 2026), and JeanGrey. Interestingly, JeanGrey is an AT-rich corynephage whose infection is restricted through interaction with the Lsr2-type protein CgpS (Badia Roige et al. 2026). A small number of additional phages capable of infecting *C. glutamicum* ATCC 13032 can be found in PhagesDB, a curated, open-access database of newly isolated bacteriophages infecting Actinomycetota (phagesdb.org, accessed October 2026) (Russell and Hatfull 2017); none of their infection dynamics, however, have yet been further characterized or published.

To date, all published work on phage characterization and phage-host interactions in *C. glutamicum* ATCC 13032 has focused on strictly virulent phages, with the sole exception of φ16, a temperate phage that was initially isolated by induction from *C. glutamicum* ATCC 21792 (Lobanova et al. 2017). This leaves a substantial gap in our understanding of how temperate phages interact with this strain. Rather than producing progeny and killing their host outright, temperate phages integrate into the bacterial chromosome, allowing the cell to survive while the phage genome is replicated alongside, and protected within, the host genome (Zhang et al. 2022). Temperate phages also represent a large, mainly uncharacterized pool of mobile genetic material capable of influencing host survival and conferring fitness benefits upon integration (Brussow et al. 2004). A well-known example is *Corynebacterium diphtheriae*, in which lysogenization by a prophage enables production of diphtheria toxin, a virulence factor that enhances the bacterium’s environmental survival (Barksdale and Arden 1974). Similar cases include Shiga toxin in *E. coli* (Schmidt 2001) and cholera toxin in *Vibrio cholerae* (Waldor and Mekalanos 1996). Collectively, these examples illustrate how the exchange of mobile genetic elements between organisms can confer selective advantages, intensifying the evolutionary pressure for optimization among phages, bacteria, and competing organisms alike.

This study aims to characterize the newly isolated phage CyranoPS, a compact temperate phage, by determining its genome sequence, annotation, and comparing genetic variation across phages from cluster ER. We also ask which host factors mediate CyranoPS infection, with a focus on mycoloylated membrane proteins, and how the immunity repressor contributes to the maintenance of the lysogenic state and to protection against re-infection. Finally, we discuss the roles of the integrase and the repressor in governing the dynamics of lysogeny and spontaneous prophage induction (SPI).

## Results and Discussion

### Isolation, morphology, and genome annotation of CyranoPS

CyranoPS phage particles were isolated from a water sample collected from the river Yvette in Saint-Forget, France. An enrichment step was performed by incubating the filtered environmental sample with *C. glutamicum* MB001 (Baumgart et al. 2013), a strain lacking the foreign genetic elements CGP1, CGP2, and CGP3, which encode several of the host’s defense mechanisms. After four rounds of plaque purification, plaque size remained heterogeneous, ranging from 0.5 to 2 mm in diameter (Figure 1A). Picking from both plaque types consistently reproduced the same variation, confirming that this heterogeneity is an intrinsic property of a single phage rather than a mixed population. Plaques were turbid and most often surrounded by a halo, independently of the media chosen for cultivation.

**Figure 1.**
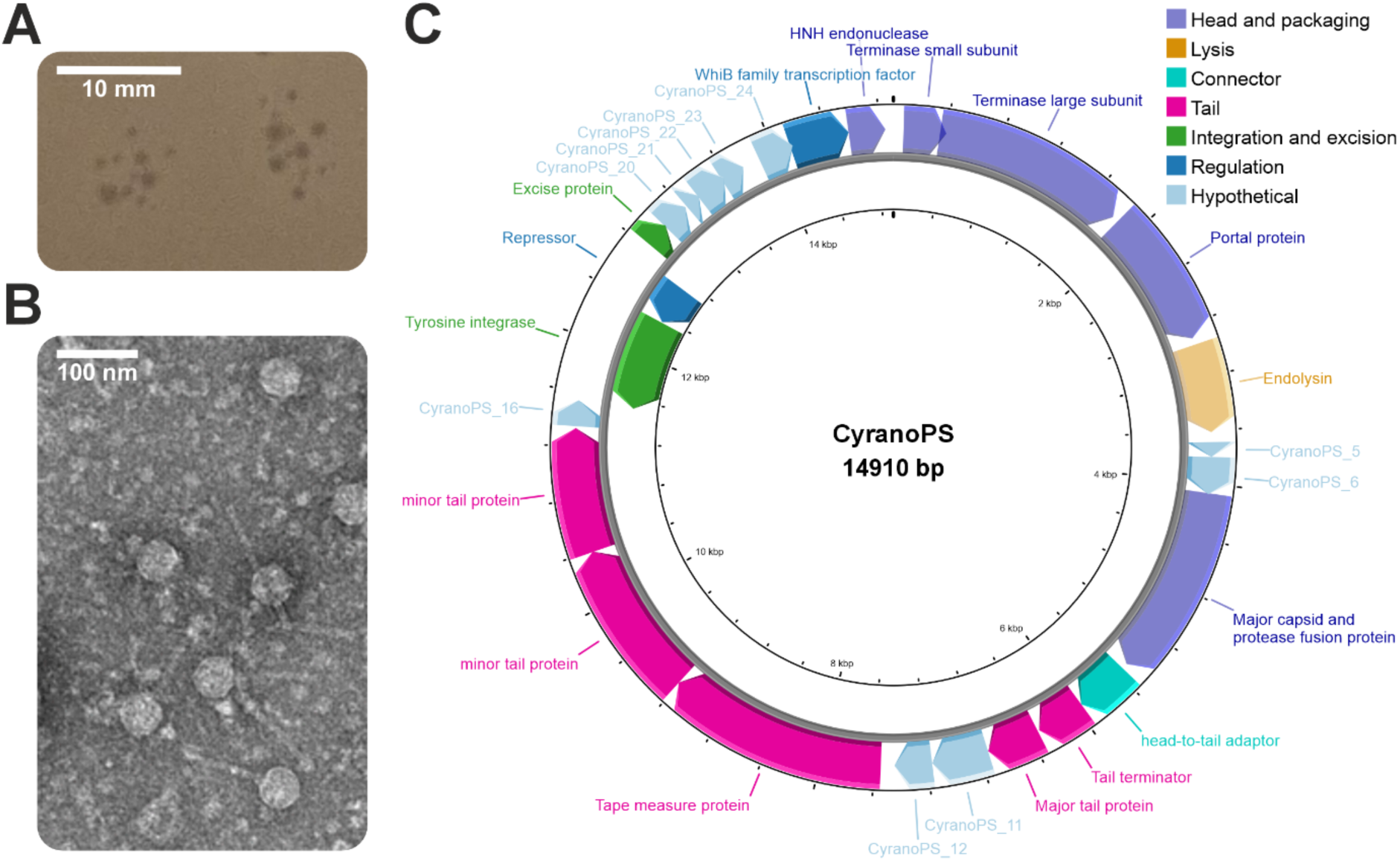
Morphology and genome organization of novel *c*orynephage CyranoPS. **(A)** CyranoPS forms heterogeneous plaques ranging from 0.5 to 2 mm in size. Scale bar, 10 mm. **(B)** TEM imaging of negatively-stained CyranoPS revealed an icosahedral capsid and an elongated non-contractile tail characteristic of a siphovirus morphology. Scale bar, 100 nm. **(C)** Annotated genome map of CyranoPS. The 26 ORFs are represented by arrows which indicate the direction of transcription. The proposed functions and categories are based on hypothetical functions predicted by bioinformatic analysis. The figure was created using Proksee (Grant et al. 2023).

Transmission electron microscopy (TEM) of filtered bacterial lysates revealed phage particles consisting of an icosahedral capsid (average diameter of 41.8 ± 1.7 nm) and an elongated, non-contractile tail (90.5 ± 1.7 nm in length), dimensions and morphology which are consistent with a siphovirus (Figure 1B).

Purified phage DNA was cleaved by several restriction enzymes. Its susceptibility to cleavage by HindIII, NspI, HaeIII and SacII suggest that CyranoPS DNA is not modified with *N*^6^-methyladenine or 5-methylcytosine (Supplementary Figure S1). Illumina sequencing revealed a compact genome of 14,910 bp with a GC content of 53.0%, close to that of the host (54.3%). The genome termini carry 14-base 3ʹ overhangs (TCCCGCGCCAGCCC), consistent with cohesive (cos) ends.

The genome was annotated and quality-checked through the SEA-PHAGES program (Science Education Alliance – Phage Hunters Advancing Genomics and Evolutionary Science), identifying 26 coding sequences and no tRNA genes (Figure 1C). Three quarters of the genome (Cyr_1–Cyr_16) carry genes likely involved in virion assembly and host lysis: terminase large and small subunits, portal protein, major capsid protein, head-to-tail adaptor, tail terminator, major tail protein, tape measure protein, and minor tail proteins along with an endolysin, all transcribed in the forward direction within a putative operon. Five genes in this region, Cyr_5, Cyr_6, Cyr_11, Cyr_12, and Cyr_16, escaped functional predictions. Gene Cyr_5, located downstream of the endolysin gene, encodes a 36-residue peptide with a predicted transmembrane domain, features that are compatible with holins.

Downstream of this structural/lysis module lies the immunity cassette characteristic of temperate phages, consistent with the lysogenic lifestyle of CyranoPS. This cassette is governed by divergent promoters: genes encoding the putative immunity repressor (Cyr_18) and tyrosine integrase (Cyr_17) are transcribed in the reverse orientation, while the excise gene (Cyr_19) and subsequent genes are transcribed in the forward orientation. Downstream of the lysogenic module, Cyr_25 and Cyr_26 encode a putative WhiB-family transcriptional regulator and HNH endonuclease, respectively. Among the five remaining genes, which lack recognizable homologs, Cyr_21 encodes a 39-residue peptide with a putative transmembrane domain.

### CyranoPS is a member of the actinophage cluster ER

Since the discovery of CyranoPS, five closely related phages were isolated by the Community Phages program (Bernhardt, unpublished) and their draft genome sequences made available in the actinobacteriophage database PhagesDB (phagesdb.org, accessed in October2026)(Russell and Hatfull 2017). Together with CyranoPS, they were grouped within a new cluster. This cluster ER comprises six phages (i.e. CyranoPS, CarefulCloud, Flashwig, HorseRider, Jalioh, and TailX7) with an average GC content of 53.0% and an average genome size of 14,989 bp (Figure 2). BLAST analysis of the encoded genes showed that the first part of the genome (encompassing the structural, assembly, and lysis genes up to the tyrosine integrase (Cyr_17), is highly conserved within the cluster. Across this region, CyranoPS shares 79-100% identity with CarefulCloud, its closest match within this region. The one exception is Cyr_15, annotated as a minor tail protein, which shares only 32% identity (Figure S2). Structural modeling of the protein encoded by Cyr_15 reveals a predicted trimeric fiber (Figure 3A), with a C-terminal end that folds like the receptor-binding protein (RBP) head domain from *Lactococcus lactis* phage bIL170 that binds polysaccharides (Figure 3B)(Ricagno et al. 2006). As the minor tail protein could be a strong candidate for mediating recognition of host receptors during adsorption (Filik et al. 2022), the divergence in its sequence within cluster ER phages suggests that although these phages belong to the same cluster, they may not all interact with the host in the same way.

**Figure 2.**
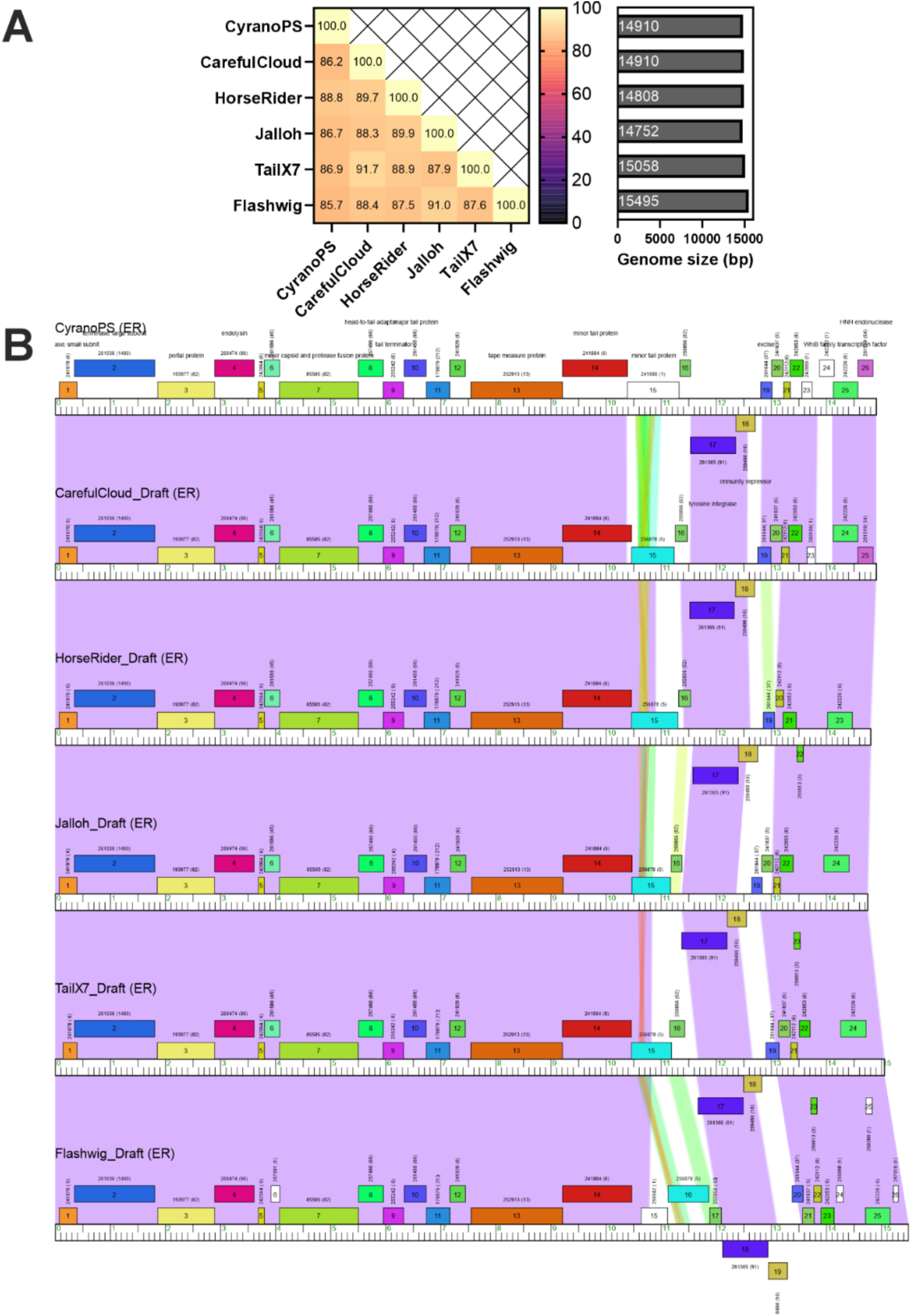
Comparison of CyranoPS to other corynephages of cluster ER. **(A)** Intergenomic similarity calculated using the tool VIRIDIC (Moraru et al. 2020) and genome sizes of cluster ER phages shows the high similarity between these related phages. **(B)** Genome alignments of CyranoPS to the cluster ER phages shows homologous genomic structures and gene functions. Alignments were made using Phamerator (https://phamerator.org, accessed in October 2026) (Cresawn et al. 2011). Open reading frames are depicted as boxes and colored according to their phamily designation (phams). Gene numbers are depicted within the boxes, and the pham number and number of pham members are shown in parantheses above each gene. Nucleotide sequence similarity is shown as colored shading between genomes, with violet being the most similar (sequence identity above 90 %) and red being the least similar. Autoannotated genomes from the Community Phages program have not been manually curated and bear the suffix *_Draft*.

**Figure 3.**
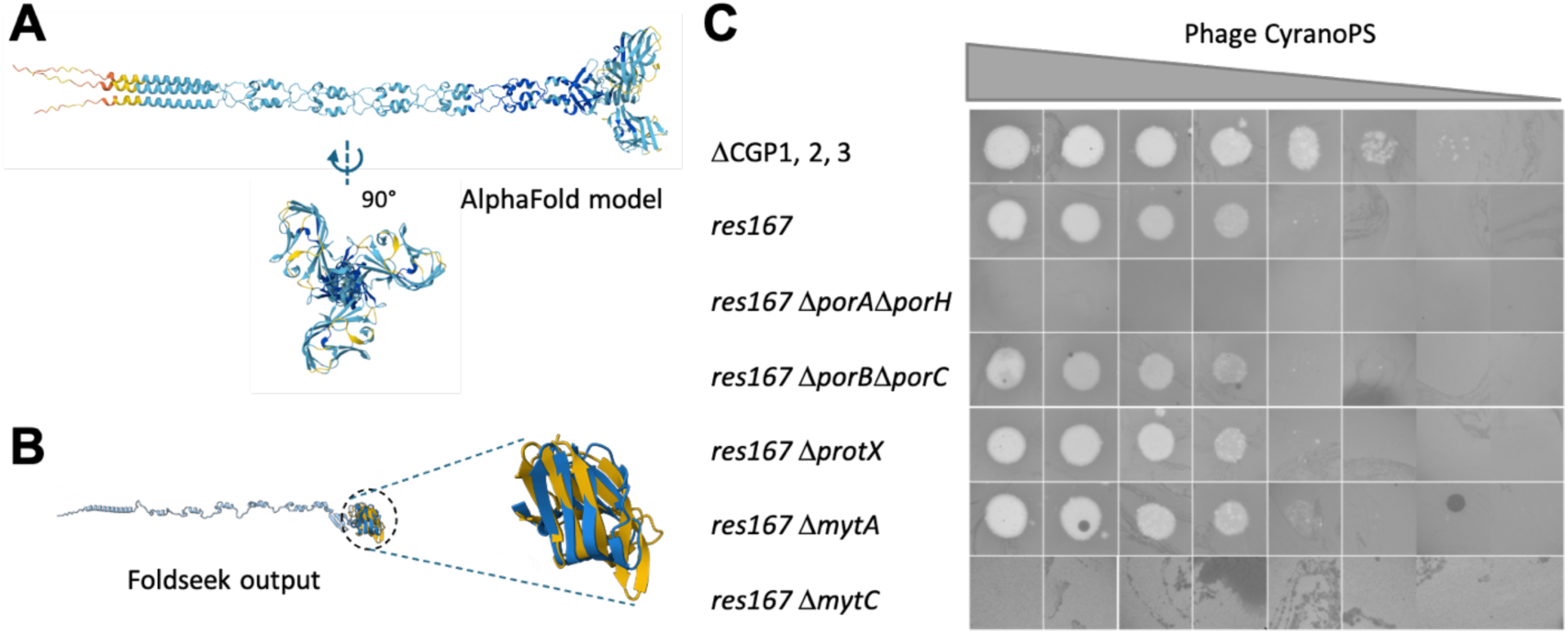
Prediction of CyranoPS receptor binding protein structure and determination of host envelope factors that facilitate infection. **(A)** Model of trimeric minor tail protein encoded by gene CyranoPS_15, predicted with AlphaFold 3 (ipTM=0.59, pTM=0.59). **(B)** Superpositions of the CyranoPS monomeric minor tail protein (in blue) with the RBP of *Lactococcus lactis* phage bIL170 (in yellow) (TM-score=0.4551; RMSD=5.14) derived from the output of the Foldseek search using a trimer as input (Probability=0.99, Seq. Id.=18.5; E-value=2.18e-2). **(C)** Impact of mutations in mycolate outer membrane proteins PorAH, PorBC, ProtX and in mycoloyltransferases MytA and MytC on the efficiency of plaquing by CyranoPS. Ten-fold serial dilutions of phage CyranoPS were prepared, and five microliters were deposited on a lawn of the indicated host strain.

The immunity repressor, encoded by Cyr_18, shows its highest similarity to the HorseRider repressor (97% amino acid identity via BLASTP in PhagesDB), compared to only 79% identity with that of CarefulCloud. Comparing the repressor sequences across all cluster ER phages, we found higher variability concentrated in the first half of the protein. This region comprises an unstructured N-terminus (residues 1–15), the predicted dimerization region (residues 15–25 in CyranoPS), and the DNA-binding HTH domain (residues 29–50 in CyranoPS). In contrast, the second half of the protein (residues 51–115) is highly conserved across the cluster (Figure S3). This variability pattern could reflect differences in how these repressors dimerize or possibly engage the transcription machinery, even as specific amino acid positions within the predicted DNA-binding region remain conserved (Q31, L34–A35, G39–R42, E49–Q50) (Figure S3).

Downstream of the immunity repressor lies a region that contains the lytic module gene *excise*, the regulator gene *whiB*, and several genes of unknown function in between. This region shows the highest variability within the cluster. Cyr_24 is the most striking case: it has no homolog in any other cluster member, and no function could be assigned to this coding region. Excise is comparatively well conserved, showing its highest similarity to HorseRider at 92% amino acid identity. Notably, the repressor, excise, and integrase (all controlled by the same divergent promoters) are the genes most conserved between CyranoPS and HorseRider, suggesting a shared regulatory network governing the lysis/lysogeny decision.

### PorAH and MytC are required for CyranoPS infection

Bacteria of the Mycobacteriales are characterized by a complex, multi-layered envelope (Jankute et al. 2015). To identify cell envelope factors that contribute to infection by CyranoPS, we tested a panel of deletion mutants in genes encoding mycomembrane proteins and mycoloyltransferases. Four pore-forming proteins have been identified in *C. glutamicum*: PorA, PorB, PorC, and PorH (Costa-Riu et al. 2003) that form hetero-oligomeric PorAH and PorBC pores, while PorA can additionally form pores on its own (Barth et al. 2010). Another mycomembrane protein is ProtX, whose only associated phenotype is its requirement for adsorption by phage Cog (McKitterick and Bernhardt 2022). Deletion of *porA* and *porH* abolished plaque-forming by CyranoPS, whereas inactivation of *protX* or deletion of *porB* and *porC* had no effect on infection (Figure 3C).

The proper localization of PorAH in the mycomembrane relies on the mycoloylation of both proteins, a post-translational modification involving mycolic acids (Carel et al. 2017; Huc et al. 2010). While MytA mediates the transfer of mycolic acid to trehalose or arabinogalactan, MytC (also called Cmt1) is the sole identified transferase in *C. glutamicum* responsible for the O-mycoloylation of proteins (Huc et al. 2010). Deletion of *mytA* did not reduce CyranoPS plaquing efficiency, whereas deletion of *mytC* completely abolished it (Figure 3C), consistent with the role of MytC in the proper localization of the porin PorAH.

Among previously characterized phages, phage CL31 and Cog both require components of the arabinogalactan and mycolic acid synthesis pathways for successful infection of *C. glutamicum*, (Hunnefeld et al. 2021; McKitterick and Bernhardt 2022). Interestingly, Cog also requires mycomembrane proteins ProtX and PorAH: while ProtX is involved in Cog adsorption, PorA is proposed to play a role after adsorption in guiding the formation of a viral channel across the envelope for DNA delivery (McKitterick and Bernhardt 2022). Given that PorA is one of the most abundant proteins in the mycomembrane, its loss may also more broadly disturb the organization of other membrane proteins that are required for infection, since its deletion lead to altered mycomembrane protein fractionation profiles compared to wild type (Marchand et al. 2012). While our work implicates PorAH in the infection by CyranoPS, structural modeling of the minor tail protein Gp15 suggests that it could bind polysaccharides. Further work is needed to test whether CyranoPS primarily adsorbs onto *C. glutamicum* cell surface polysaccharides, and that subsequent steps of infection require the host porins PorAH.

Taken together, our analysis of CyranoPS broadens our understanding of how phages exploit *C. glutamicum* known host cell surface proteins for infection: relying on both ProtX and PorAH like phage Cog, PorAH only like CyranoPS, or requiring none of them like CL31.

### CyranoPS integrates into the host genome to form stable lysogens

Infection of bacterial lawns with CyranoPS consistently produced regrowth within the clearing zone after 2-3 days, indicating survival of a subpopulation of cells. Colonies recovered from the regrowth zone were purified through multiple rounds of restreaking, yielding strains with a stable phenotype over successive passages, which can be a first indication of true lysogeny rather than transient resistance. Streaking these putative lysogens onto a lawn of the parental, phage-sensitive *C. glutamicum* MB001 produced a clear lysis zone in every case (Figure 4A), indicating that the isolated strains actively carry and release infectious phages, consistent with a stable lysogenic state. Notably, despite harboring an inducible prophage (with the associated risk of SPI and cell death) the lysogens showed no significant growth defect in monocultures relative to the non-lysogenic parental strain (Figure 4B), indicating that CyranoPS carriage imposes little to no fitness cost under standard culture conditions.

**Figure 4.**
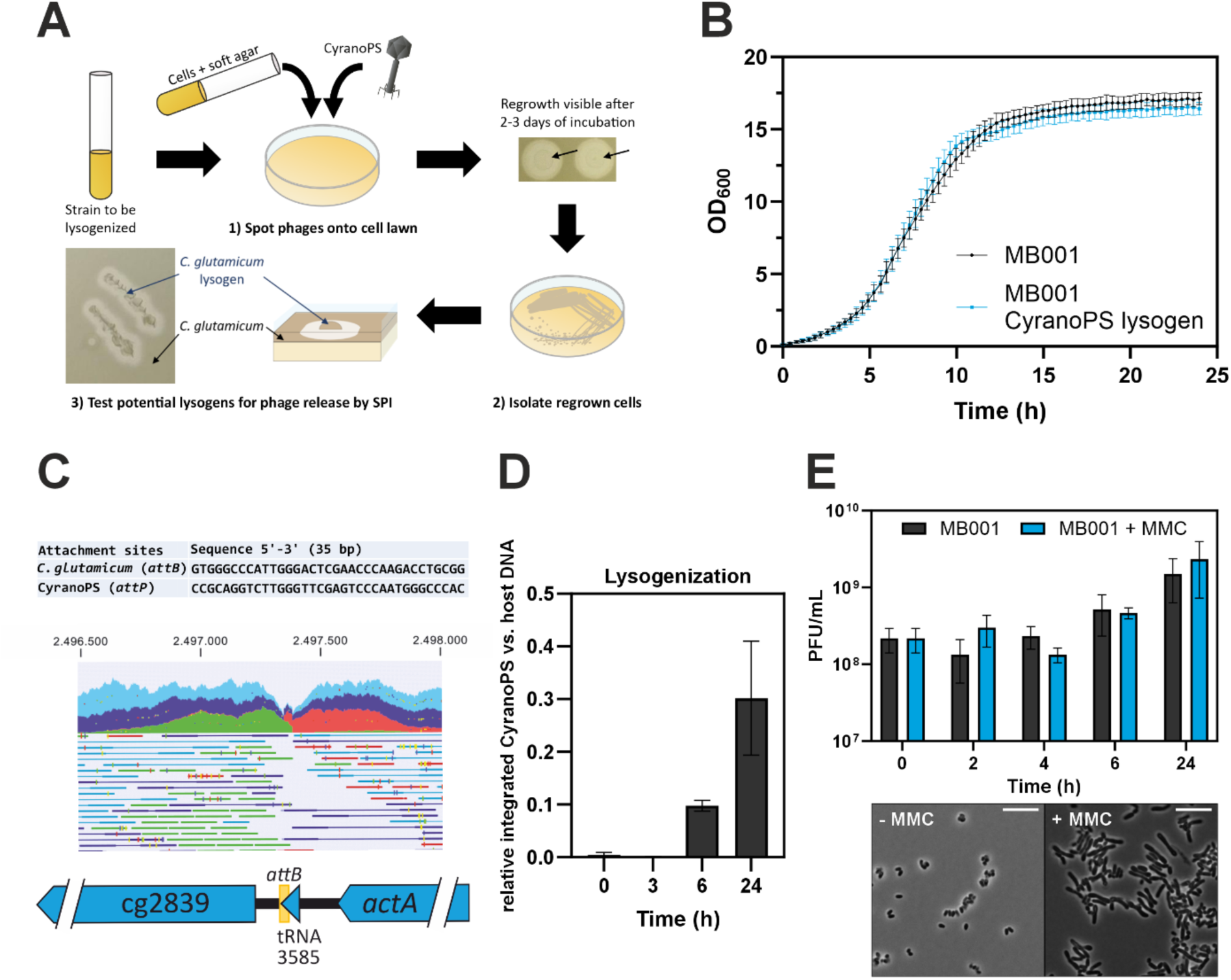
Generation and characterization of *C. glutamicum* MB001 CyranoPS lysogens. **(A)** Workflow to generate CyranoPS lysogens using the prophage-free strain *C. glutamicum* MB001. Successful genomic integration was verified by means of spontaneous induction (SPI) and lysis of the phage-susceptible strain MB001 in the overlay agar. **(B)** Growth of *C. glutamicum* MB001 compared to the lysogen in LB containing 5 mM CaCl_2_ (n=3). **(C)** Sequence of the attachment sites of the CyranoPS (*attP*) and *C. glutamicum* (*attB*) genome. Genome sequencing of isolated lysogens confirmed the position of chromosomal integration within a gene encoding a lysine tRNA (cgtRNA_3585). Mapped reads displayed in CLC Genomics Workbench show disruption of paired reads due to prophage integration at *attB* (green: single forward direction; red: single reverse direction; light blue: paired-end reads; dark blue: broken paired reads; yellow: ambiguously mapped). **(D)** Kinetics of lysogenization measured by qPCR revealed an increase of CyranoPS integration over time. **(E)** CyranoPS lysogens show a high level of spontaneous induction, which is not further increased upon SOS induction by mitomycin C (MMC, 0.6 µM) (n=3). Light microscopy revealed elongation of cells treated with MMC, confirming that treated cells underwent the expected stress response. Scale bar, 10 µm in length.

Integration requires a site of homology between the phage and bacterial genomes. CyranoPS encodes an armless tyrosine integrase that belongs to the same class as that of the well-characterized mycobacteriophage Brujita (with a 32.56% identity over a query coverage of 90% per BLAST analysis), which has been reported to require a 39-bp homology sequence for integration (Lunt and Hatfull 2016). Based on this, we screened the CyranoPS and *C. glutamicum* strain MB001 genomes for candidate attachment sites (*attP*/*attB*). This search identified two candidates: the first sites (20 bp) are localized to the positions 4201 (*attP*) and 1,369,413 (*attB*) in the viral and bacterial genomes, respectively. The second 35-bp sites locate to the position 12,409 for CyranoPS *attP* and position 2,497,382 for *C. glutamicum* MB001 *attB* on the complementary strand. The latter *attP* is located within the immunity repressor gene (Cyr_18) near its 3’ end. To distinguish between these candidates, we sequenced the isolated lysogenic strains and mapped the Illumina reads against the parental genome. This revealed an assembly break precisely at the 35-bp site, which disrupts a host tRNA gene (Lys-tRNA-3585), pointing to this site as the main attachment site used by CyranoPS (Figure 4C). qPCR with a corresponding primer set confirmed integration at this location. Infections in liquid culture showed detectable integration within 6 hours, with approximately 10% of the cells carrying the prophage, rising to 30%± 10% by 24 hours (Figure 4D).

### Induction of CyranoPS lysogens is SOS-independent

Prophage induction in a population of lysogenic cells can be regulated through several mechanisms. The most extensively characterized lytic-lysogenic switch is that of the *E. coli* phage λ, which depends on the SOS-response: the lysogenic state is maintained by the CI repressor through inhibition of the early promoters pL and pR (Brady et al. 2021a). The SOS response (activated by DNA damage) triggers RecA activation, which promotes auto-proteolysis of LexA; the same RecA-ssDNA filaments also drive self-cleavage of the CI repressor via a LexA-like autoproteolytic mechanism, directly linking prophage induction to the host DNA damage response (Zhang et al. 2022). Although addition of mitomycin C (MMC) triggers the SOS response through DNA damage, it did not increase induction rates of CyranoPS over the course of cultivation (Figure 4E), raising the question of whether CyranoPS instead follows an SOS-independent induction mechanism, as described for other phages such as phi3T or φ105 in *B. subtilis* (Brady et al. 2021b; Erez et al. 2017). Spontaneous prophage induction of CyranoPS resulted in a high titer of 10^9^ PFU/mL after 24 hours of cultivation. This is among the highest values reported for a prophage. A similar titer was described for the *Salmonella* Typhimurium prophage BTP1, whereas most other prophages release between 10^2^ and 10^7^ PFU/mL without induction (Owen et al. 2017). As stated above, the mycobacteriophage Brujita also encodes an armless tyrosine integrase and shows high SPI (Broussard et al. 2013). The lack of directionality in recombination by armless integrases could contribute to this phenotype (Lunt and Hatfull 2016). The Brujita repressor, however, shares only 25.4% global amino acid identity with Cyr_18, so repressor similarity is unlikely to explain the high SPI of both phages.

### Integration-dependent truncation of Cyr_18 controls lysogeny and superinfection immunity

During generation of lysogenic *C. glutamicum* strains with CyranoPS, we found that *attP* is located within the 3’ end of the immunity repressor gene, Cyr_18. Upon integration of the phage DNA into the bacterial chromosome (*attB*), the encoded repressor protein lacks its C-terminal 15 residues, and the integrase gene simultaneously loses its promoter (Figure 5A and 5B). As the repressor plays a central role in the lysis/lysogeny switch, we hypothesized that it would also confer immunity against lytic infection by incoming, free CyranoPS particles.

**Figure 5:**
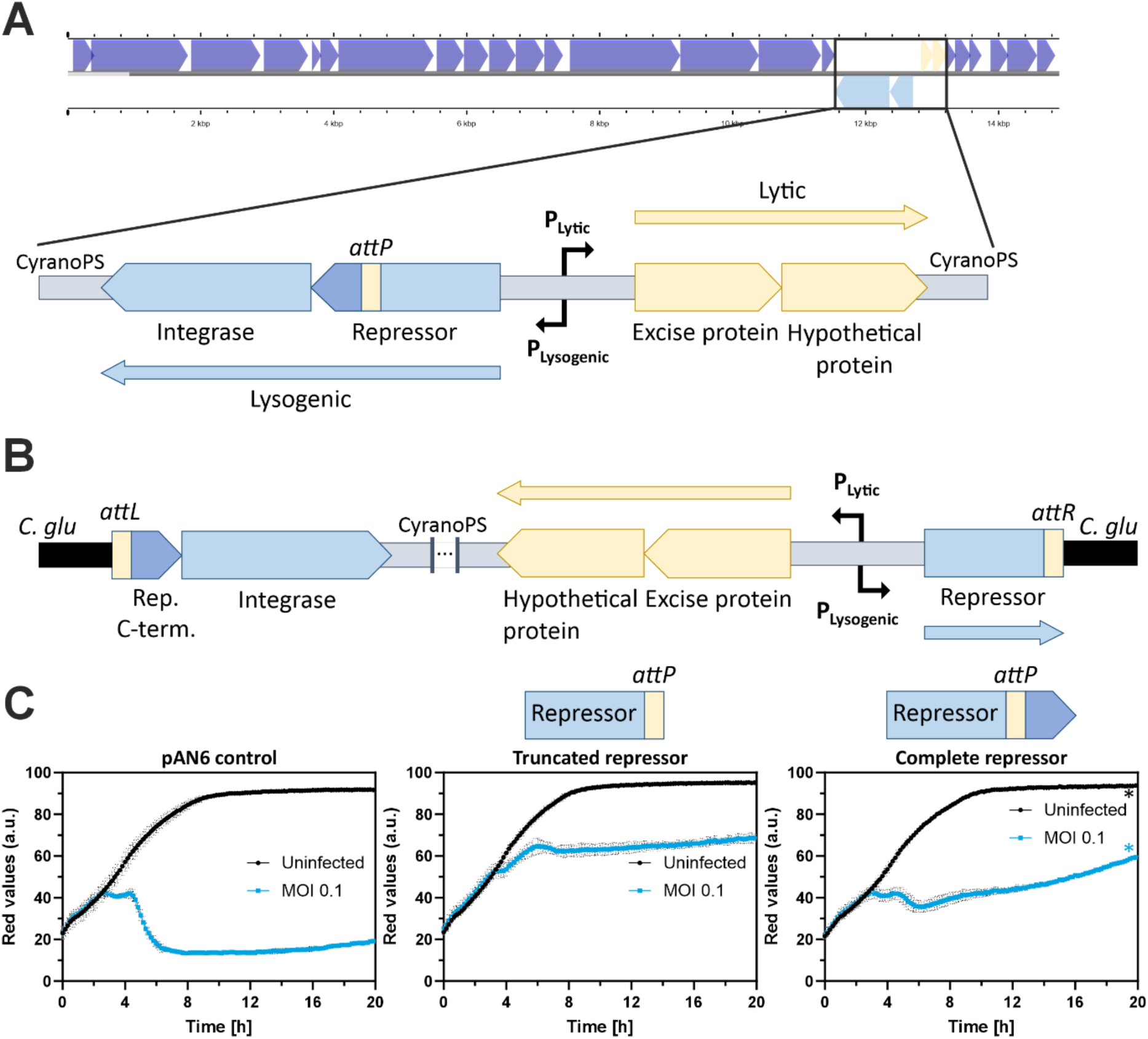
The lysis-lysogeny switch of CyranoPS depends on an integration-dependent truncation of the central phage repressor (Cyr_18). **(A and B)** The CyranoPS genome map generated on Proksee (Grant et al. 2023). Shown are the lytic (A) and integrated (B) versions of the lytic/lysogenic cassette of CyranoPS, which is under the transcriptional control of the putative promoters P_Lytic_ and P_Lysogenic_. Gene sizes are not drawn to scale. Upon lysogeny, integration of the CyranoPS genome within the bacterial chromosome leads to the production of a truncated repressor missing the C-terminal 15 residues. **(C)** Growth curves of prophage-free *C. glutamicum* MB001 strain expressing plasmid-encoded repressor variants and infected with CyranoPS upon inoculation (n = 3 biological replicates; * n = 2). Superinfection immunity conferred by repressor expression is used to determine repressor activity.

To test this, we generated two overexpression plasmid systems: one encoding the full-length repressor, corresponding to its form prior to genome integration, and one encoding the truncated repressor, corresponding to its form within the CyranoPS prophage. *C. glutamicum* MB001 cells overexpressing each version were then challenged with CyranoPS infection (MOI 0.1). Cells expressing the truncated repressor showed a marked reduction in infection, indicating that this truncated form confers immunity against superinfection. In contrast, cells expressing the full-length repressor showed an intermediate phenotype, though cultures still displayed visible recovery and regrowth (Figure 5C). This suggests that the truncated repressor is sufficient to block the expression of the lytic cassette of the incoming phages. In a lysogen, the occupied *attB* site by the resident CyranoPS prophage would additionally be expected to prevent the integration of the infecting phage genome. Nevertheless, this second effect was not tested upon repressor expression in non-lysogenic cells. This infection event by a second phage would, however, still lead to expression of the full-length repressor from the incoming genome, and both proteins could accumulate within the cell.

In the mycobacteriophage Brujita, which shares the same integrase class and repressor system as CyranoPS, the C-terminus of the full-length repressor has been described as a destabilizing tag targeted by the host protease systems (Broussard et al. 2013), limiting the accumulation of the full-length protein. Furthermore, accumulated integrase levels may reach sufficient levels to trigger SPI of the resident prophage (Broussard et al. 2013). Notably, the CyranoPS immunity cassette encodes an Excise protein sharing an identity of 40.43% with the Cro-like regulator of the Brujita system, though it remains unclear how this protein affects the switch. Further work will be needed to determine whether CyranoPS follows a similar regulatory mechanism.

## Conclusion

CyranoPS is the first temperate corynephage isolated on the prophage-free strain *C. glutamicum* MB001, which is derived from the industrial strain ATCC 13032 (Baumgart et al. 2013). Following its isolation from river water in France, five related phages were isolated in the USA with gene content similarity above 35% with CyranoPS (Phagesdb.org) (Russell and Hatfull 2017), compelling the creation of the new phage cluster ER. To our knowledge, phages of this cluster have the smallest genomes (14.8 to 15.5 kb) of all *Corynebacterium* phages described so far.

Our data suggested that CyranoPS infection requires MytC-mediated protein mycoloylation and potentially the PorAH porin. Notably, phages CyranoPS and Cog both depend on PorAH, whereas CL31 does not (McKitterick and Bernhardt 2022). These phages show how diverse phages exploit (or are restricted by) this shared bacterial architecture. However, whether surface polysaccharides serve as the primary receptor (as suggested by structural similarity between its minor tail protein and a Lactococcal phage RBP (Ricagno et al. 2006)) remains to be determined.

We demonstrated that CyranoPS lysogenizes *C. glutamicum*. About 10% of the cells carried the prophage after 6 hours, and 30% ± 10% after 24 hours. Notably, chromosomal integration of CyranoPS truncates the immunity repressor Cyr_18 by 15 C-terminal amino acids, with the truncated form providing stronger protection against superinfection than the full-length protein. Together with the high rate of spontaneous prophage induction and the lack of enhanced induction upon SOS activation, this suggests a potentially SOS-independent lytic–lysogenic switch. However, its underlying regulation, including the role of Excise, remains to be resolved, as in related phage systems (Broussard et al. 2013). More broadly, comparative analysis and functional characterization of related phages from the newly established cluster ER may help to unravel the molecular mechanisms governing their lytic-lysogenic switch and provide new insights into the regulation of lysogeny in compact temperate actinobacteriophages.

## Methods

### Bacterial strains and growth conditions

All bacterial strains used in this study are listed in Supplementary Table S1. *Escherichia coli* and *C. glutamicum* strains were cultivated in Lysogeny Broth medium (Miller LB: 10 g L⁻¹ tryptone, 5 g L⁻¹ yeast extract, 10 g L⁻¹ NaCl) at 37 °C and 30 °C, respectively. When required, kanamycin was added at a final concentration of 25 µg mL⁻¹. Liquid cultures were inoculated from single colonies obtained from fresh 1.5% (w/v) BHI agar plates (37 g L⁻¹ BHI; Difco, BD, Heidelberg, Germany) or LB agar plates and incubated for up to 18 h at 30 °C. Pre-cultures were grown in 1 mL medium in 2.2 mL square V-bottom 96-deep-well plates (VWR, Radnor, PA, USA) sealed with breathable rayon film (VWR) and incubated at 30 °C and 900 rpm in a Microtron incubator shaker (Infors, Bottmingen, Switzerland). For larger cultures, cells were grown in 5 mL medium in culture tubes at 170 rpm and 30 °C.

### Recombinant DNA work and chromosomal deletions

Overexpression constructs for the repressor gene (Cyr_18) were generated using plasmid pAN6 (Frunzke et al. 2008b), which contains a *Ptac*-controlled expression cassette with a *lacI* repressor. Target genes were amplified by PCR, assembled into the vector backbone using Gibson assembly, and transformed into *E. coli* DH5α by heat shock. Plasmids generated and primers used in this study are listed in Supplementary Tables S2 and S3, respectively. Plasmids were verified by sequencing and subsequently introduced into *C. glutamicum* by electroporation (Bio-Rad; 2.5 kV, 25 µF and 200 Ω). Expression of recombinant proteins was induced by addition of IPTG at a final concentration of 100 μM.

### Phage isolation, plaque determination, and propagation

River water was sampled in Saint Forget, France (GPS coordinates: 48.70192 N, 2.021751 E). The sample was first passed through a 0.22-µm filter and then supplemented with Lennox LB (10 g L⁻¹ tryptone, 5 g L⁻¹ yeast extract, 5 g L⁻¹ NaCl), 1 mM CaCl_2_, and *C. glutamicum* strain MB001. After incubation for 24 h at 30°C, the filtered supernatant was spotted on a double agar layer in the presence of MB001 and incubated for 24 h at 30°C. Plaques were purified through four rounds of picking and plating.

For double-agar overlay assays, *C. glutamicum* precultures grown in Miller LB medium containing 5 mM of CaCl_2_ were mixed with phages and LB soft agar (0.4% agar) to a final OD₆₀₀ of 0.3. The mixture was poured onto LB agar plates and allowed to solidify. For spot assays, tenfold serial dilutions of phage lysates prepared in SM buffer (50 mM Tris–HCl pH 7.5, 8 mM MgSO₄, 100 mM NaCl) were spotted onto the surface of solidified soft agar containing bacteria and CaCl_2_. Plates were incubated at 30 °C for 18–24 h before plaque formation was assessed. The phage titer could be determined by counting single plaques and calculating the plaque forming units (PFU) per mL.

### Microscopy

For observation of phage particles by negative staining TEM, CyranoPS phages were collected after flooding highly confluent double-agar overlay plates with phage buffer (10 mM Tris-HCl pH 7.5, 10 mM MgSO_4_, 1 mM CaCl_2_, 68 mM NaCl) for 24 hours at 4 °C and subsequent passage through a 0.22-µm filter. Phages were stained with 2% wt/vol uranyl acetate and observed with a Tecnai Spirit microscope operated at 120 kV (TFS) and equipped with a Rio9 camera (Ametek/Gatan). The magnification used was 15,000× with a pixel size of 0.44 nm at the level of the specimen.

For light microscopy, bacteria were observed at 100x magnification in oil immersion using a Zeiss Axio Imager.M2 microscope (Zeiss, Jena, Germany). For imaging, 1.5 μL of the respective culture was pipetted onto an agarose pad prepared with 1% agarose. Images were acquired with an AxioCam ICc 3 camera (Zeiss, Jena, Germany) and AxioVision software (Zeiss, Jena, Germany).

### DNA sequencing and genome assembly of phages and lysogenic strains

Phage DNA was extracted from a high titer lysate using the PCI/SDS protocol (https://phagesdb.org/media/workflow/protocols/pdfs/PCI_SDS_DNA_Extraction_2.2013.pdf). Illumina whole genome sequencing of the phage DNA was performed at the NGS platform of the I2BC. NextSeq 500/550 High Output Kit v2 was used to prepare the library that was sequenced using Illumina NextSeq NB552053, yielding 12,261,187 paired-end 75-base reads. Raw reads were trimmed with cutadapt 3.2 and filtered with Skewer 0.2.2 prior to assembly (Jiang et al. 2014; Martin 2011). De novo assembly was performed with Newbler v2.9 as previously described (Russell 2018) and further checked with Consed v.29 and Unicycler (Gordon et al. 1998; Wick et al. 2017).

Genomic DNA was isolated from the bacterial lysogens using the NucleoSpin Microbial DNA Mini Kit (Macherey-Nagel, Düren, Germany). Sequencing libraries were prepared with the TruSeq DNA PCR-Free Sample Preparation Kit (Illumina, Chesterford, UK) according to the manufacturer’s instructions. The libraries were quantified with the KAPA Library Quantification Kit (Peqlab, Bonn, Germany) and normalized before pooling. Paired-end sequencing (2 × 150 bp) of the pooled libraries was carried out on an Illumina MiSeq platform. Base calling and initial processing were done with the Illumina instrument software, which generated FASTQ files for downstream analysis.

Further data processing followed a workflow adapted from (Keppel et al. 2020). Adapter sequences were first trimmed from the reads. To reduce PCR amplification artifacts, reads were then collapsed for each sample prior to mapping. The processed reads were mapped to the *C. glutamicum* MB001 reference genome (NCBI accession CP005959) using CLC Genomics Workbench v24 (Qiagen, Hilden, Germany). Mapping was performed with a length fraction of 0.9 and a similarity fraction of 0.9. An interruption of the alignment marked the phage integration site. At this position, the phage sequence was added manually in its integrated form.

### Structural and functional genome annotations and comparisons

Structural annotation was performed using Glimmer (Delcher et al. 2007) and GeneMark v2.5p (Besemer and Borodovsky 2005), PHANOTATE (McNair et al. 2019), Aragorn v1.2.40 (Laslett and Canback 2004), tRNAscan-SE v2.0 (Chan et al. 2021), and Phamerator (Cresawn et al. 2011). Start sites were selected based on gene length, minimal gaps or overlaps, RBS scores, and BLASTP alignment.

Functional annotation was performed with BLASTP (Altschul and Lipman 1990) using PhagesDB and NCBI non-redundant databases (e values < 0.001), HHpred (Soding et al. 2005) using databases PDB_mmCIF70_30_Mar, Pfam-A_v37, Uniprot-Swissprot-viral70_3_Nov_2021, and NCBIConservedDomain(CD)_v3.19, and PHOLD (Bouras et al. 2026). Protein structural prediction was performed using AlphaFold3 (Abramson et al. 2024) and followed by searches for homologous structures using Foldseek (van Kempen et al. 2024). Membrane proteins were predicted with DeepTMHMM v1.0.42 (Hallgren et al. 2022).

Genome comparisons were performed using Phamerator (Cresawn et al. 2011), VIRIDIC (Moraru et al. 2020) and CAGECAT clinker analysis (Gilchrist and Chooi 2021). Selective protein alignments (were carried out using CLUSTAL W and visualized using Jalview 2 (Thompson et al. 1994; Waterhouse et al. 2009).

### Isolation of bacterial lysogens and determination of lysogenization kinetics, superinfection immunity and spontaneous induction

Lysogens were isolated from the regrowth that appeared within the clearing zone of CyranoPS-infected bacterial lawns (decimal dilution spots) after 2–3 days of incubation at 30 °C. Individual colonies were picked from this zone and purified by three rounds of restreaking on LB agar plates. To test whether the purified strains released phage particles, they were streaked onto a lawn of the parental, phage-sensitive *C. glutamicum* MB001. Plates were incubated at 30 °C overnight and inspected for lysis zones along the streak.

To test lysogenization over time, liquid cultures of *C. glutamicum* MB001 were grown in LB medium supplemented with 5 mM CaCl₂ to an OD₆₀₀ of 0.3 at 30 °C and 110 rpm. CyranoPS phage stock was added at a multiplicity of infection (MOI) of 0.3, and the cultures were incubated further under the same conditions.

For integrated phage DNA quantification, 500 µL culture samples were harvested at each time point (0 h, 3 h, 6 h, and 24 h) and washed twice with cold 1× PBS to remove extracellular phages. The cells were disrupted twice for 30 s at 6,500 rpm in a Precellys cell disruptor (Bertin Technologies, Montigny-le-Bretonneux, France). The lysates were centrifuged (14000 x g, 10 min, 4 °C), and the DNA in the supernatant was used for downstream quantification by qPCR. Phage DNA levels were quantified by qPCR using Luna® Universal qPCR Master Mix (New England Biolabs, Ipswich, MA, USA). Reactions contained 50 ng total DNA and were performed on a qTOWER 2.2 system (Analytik Jena, Jena, Germany) according to the manufacturer’s instructions. Phage DNA was detected using primers B119/B120, whereas host genomic DNA was quantified using primers B121/B122 targeting the *ddh* reference gene. qPCR reactions were performed with three biological replicates and two technical replicates per sample. Standards were generated using a dilution series of purified PCR products together with a no-template control to define the limit of detection for subsequent calculations. Relative phage DNA abundance was calculated using the comparative analysis function in qPCRsoft 3.1 (Analytik Jena), expressing phage DNA levels relative to host genomic DNA.

Induction of the CyranoPS prophage was observed to assess whether CyranoPS lysogens show increased prophage induction in the presence of the SOS-response inducing agent mitomycin C (MMC). Cells from precultures were harvested by centrifugation and washed in SM buffer twice before transferring to a new collection tube and three additional wash steps in LB supplemented with 5 mM CaCl_2_. Cultures were inoculated to an OD_600_ of 0.5 in the presence or absence of 0.6 µM MMC. Spontaneous induction of CyranoPS in lysogens was assessed by double-agar overlay assays at different time points (0 h, 2 h, 4 h, 6h and 24 h) as described above using supernatant of lysogens collected at different time points.

### Phage infection curves

Growth and infection assays were primarily monitored using the Growth Profiler 960 (EnzyScreen, Netherlands). Images of the clear plate bottom were acquired during incubation and analyzed using GP960Viewer software (v1.1.1.0). Growth curves were calculated based on the Red Value signal, which quantifies the intensity of red pixels within a defined area of each well image.

Infection assays were performed in Miller LB medium containing 5 mM CaCl_2_ with an initial OD₆₀₀ of 0.3 at 30 °C. Cultures were grown in volumes of 200 µL or 2 mL in the Growth Profiler system (96-well and 24-well, square white plates; 250 rpm; 85% humidity).

Data visualization and statistical analyses were performed using GraphPad Prism v8.0 (GraphPad Software, La Jolla, CA, USA). Graphs represent mean values with standard deviations, and statistical significance was determined using two-tailed unpaired t-tests. If not stated otherwise, n = 3.

## Data

ENA for sequencing of lysogens: PRJEB126091

Genbank: PV915806.1

## Acknowledgements

We thank Thomas G Bernhardt and the Community Phages program for sharing the unpublished sequences of their phages. We acknowledge the High throughput sequencing facility of l2BC for its sequencing and bioinformatics expertise, Daniel Russel for performing the assembly of the CyranoPS genome, and Véronique Delesalle for advice on genome annotation. We also acknowledge members of the Bayan team for their kind gift of *Corynebacterium* strains, Ana Arteni, Laura Pieri and Malika Ouldali for their help at the I2BC electron microscopy platform, Stéphane Roche for advice on protein structure analysis, and students, staff and faculty of the University Paris-Saclay who are key to the Phage Discovery and Phage DNA Explorer teaching units: Maylis Fabregues, Raphael Bourgeois, Albert Barkim, Ameline Batsché, Clément Cousin, Lucia Lanteri-Minet, Baptiste Ortheau Alberti, Carlyne Samson, Monique Auberdiac, Daniel Delaruelle, Cécile Labarre, Florence Constantinesco-Becker and Anne Lopes.

