## Supplementary Data for "CyranoPS, a novel compact temperate corynephage infecting *Corynebacterium glutamicum* MB001"


### Overview:

Supplementary Fig. S1: Restriction profile of CyranoPS.
Supplementary Fig. S2: Alignment of the minor tail proteins within the cluster ER corynephages.
Supplementary Fig. S3: Alignment of repressor proteins within the cluster ER corynephages.
Supplementary Table S1: Strains used or generated in this study.
Supplementary Table S2: Plasmids used or generated in this study.
Supplementary Table S3: Oligonucleotides used or generated in this study.

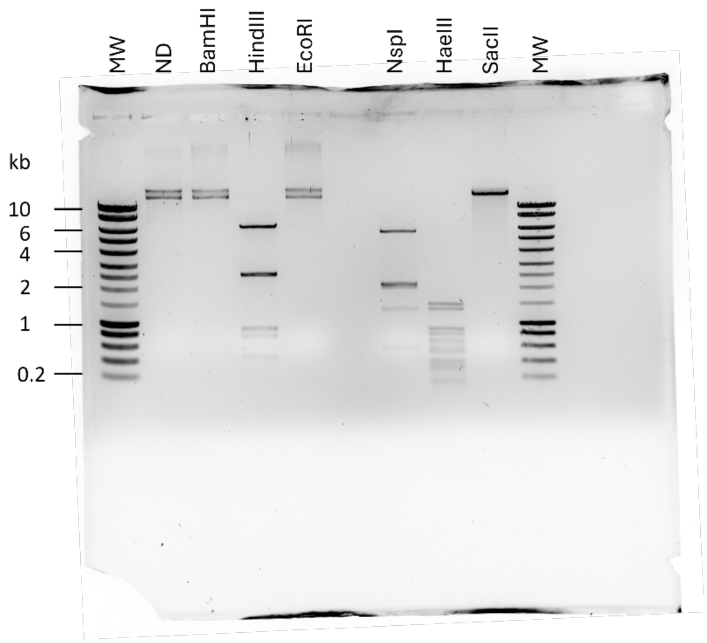


**Supplementary Figure S1. Restriction profile of CyranoPS.** Purified genomic phage DNA was incubated with several restriction enzymes (BamHI, HindIII, EcoRI, NspI, HaeIII, SacII). The molecular weight DNA standard (MW) used is a Eurogentec SmartLadder (Eurogentec, Seraing, Belgium).

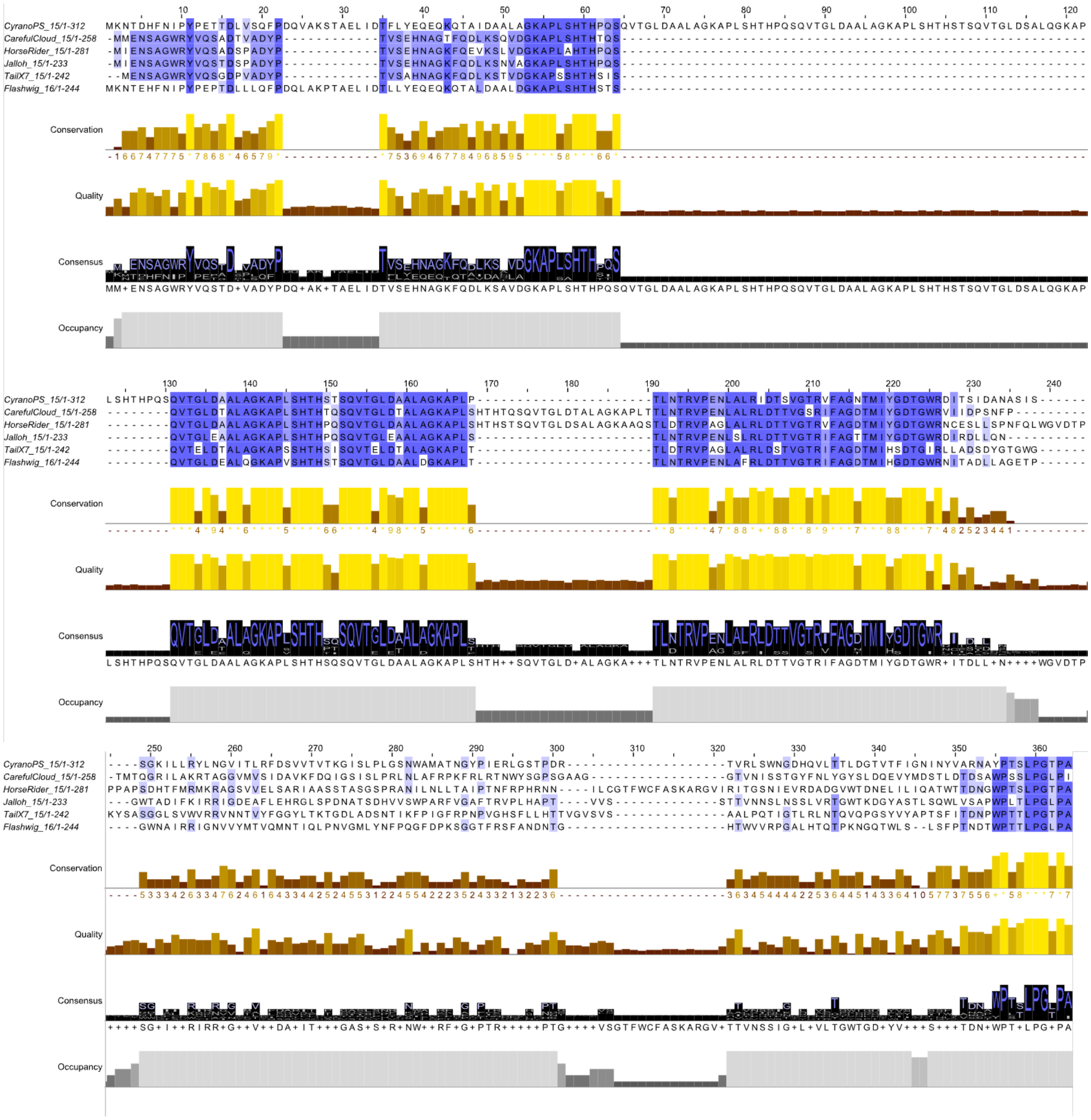

**Supplementary Figure S2. Alignment of the minor tail proteins within the cluster ER corynephages.** Unlike other structural genes of CyranoPS, these alignments reveal large gaps and unaligned regions between the minor tail proteins of cluster ER phages. Alignments were made with CLUSTAL W and visualized using Jalview 2 (Waterhouse *et al.* 2009).

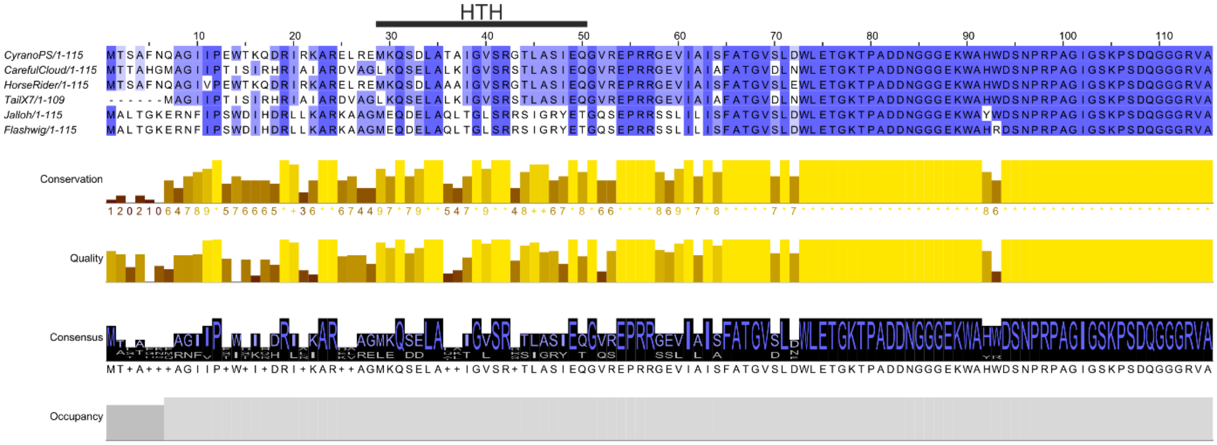


**Supplementary Figure S3. Alignment of repressor proteins within the cluster ER corynephages.** These alignments reveal higher variability in the N-terminal half of the proteins. This region comprises an unstructured N-terminus (residues 1-15), a putative dimerization region (residues 15-25) as well as the DNA-binding domain (HTH, residues 29-50). Residues 51 to 115, including the C-terminus, are highly conserved within the compared phages. Residue numbers are based on the CyranoPS sequence. Alignments were made with CLUSTAL W and visualized using Jalview 2.

### Supplemental Table S1. Strains generated or used in this study.

| Organism | Strains | Characteristics | Reference |
|---|---|---|---|
| *E. coli* | DH5α | F- Φ80*lacZ*Δ*M15* Δ(*lacZYA-argF*) *U169 recA1 endA1 hsdR17* ($r_k^-$, $m_k^+$) *phoA supE44* thi-1 *gyrA96 relA1* $\lambda^-$ | Invitrogen (Thermo Fisher Scientific, DE) |
| *C. glutamicum* | ATCC 13032 MB001 (ΔCGP1, 2, 3) | Biotin-auxotrophic strain with deleted prophage clusters CGP1 (cg1507-cg1524), CGP2 (cg1746-cg1752), and CGP3 (cg1890- cg2071) | (Baumgart *et al.* 2013) |
| *C. glutamicum* | ATCC 13032 *res167* | Restriction-deficient mutant of ATCC 13032, carries full deletions of *ttnp14a*, *cg1982-cg2033* (full region), and *cg2060*; truncations of *cg1981*, *cg2034*, *cg4005*, and *psp3*. | (Tauch *et al.* 2002) |
| *C. glutamicum* | ATCC 13032 *res167* Δ*porA* Δ*porH* | ATCC 13032 *res167* with deletion of *porA* and *porH* | (Schiffler *et al.* 2007) |
| *C. glutamicum* | ATCC 13032 *res167* Δ*porB* Δ*porC* | ATCC 13032 *res167* with deletion of *porB* and *porC* | Unpublished, a kind gift by Nicolas Bayan |
| *C. glutamicum* | ATCC 13032 *res167* Δ*protX* | ATCC 13032 *res167* with deletion of *protX* | (Issa *et al.* 2017) |
| *C. glutamicum* | ATCC 13032 *res167* Δ*mytA* | ATCC 13032 *res167* with deletion of *mytA* | (Dietrich *et al.* 2020) |
| *C. glutamicum* | ATCC 13032 *res167* Δ*mytC* | ATCC 13032 *res167* with deletion of *mytC* | (Huc *et al.* 2013) |

### Supplemental Table S2. Plasmids generated or used in this study.

| Plasmid | Characteristics | Reference |
|---|---|---|
| pAN6 | $Kan^R$; *C. glutamicum*/*E. coli* shuttle vector for gene expression under control of the *tac* promoter; ($P_{tac}$, lacIq, pBL1 oriVC.g., pUC18 oriVE.c.) | (Frunzke *et al.* 2008) |
| pAN6-CyrPS-Rep-truncated | Derivative of pAN6 containing the CyranoPS truncated repressor gene expressed during lysogeny | This study |
| pAN6-CyrPS-Rep-complete | Derivative of pAN6 containing the CyranoPS complete repressor gene expressed during the lytic cycle prior to integration | This study |

**Supplemental Table S3. Primers generated or used in this study.**

| Internal name | Sequence | Purpose |
|---|---|---|
| B398_pAN6-17_fw | TACATATGACCTGAGCTAGCATGACTAGTGCATTTAATCAAGCC | Generation of vector for repressor overexpression |
| B399_pAN6-17.1_rv (full) | AAACGACGGCCAGTGAATTCTTAGGCTACTCTGCCGCC | Generation of vector for complete repressor overexpression |
| S010_pAN6-17-trun_rv | AAACGACGGCCAGTGAATTCTTAATCCGCAGGTCTTGGGTT | Generation of vector for truncated repressor overexpression |
| pEKEx2-fw | GATATGACCATGATTACGCC | Sequencing of overexpression plasmid |
| pEKEx2-rv | CGGCGTTTCACTTCTGAGTTCGGC | Sequencing of overexpression plasmid |
| B119_qPCR_IntCyr_for | GGAGCATTCTCCAAAACAGG | Quantification of integrated phage DNA by qPCR |
| B120_qPCR_IntCyr_rev | CACCCCCGAACATGAGATAG | Quantification of integrated phage DNA by qPCR |
| B121_qPCR_ddh_fw | CAACCACACCGTGGAATACA | Quantification of host DNA by qPCR |
| B122_qPCR_ddh_rev | TTCTCTGGGGAGAGCAGGTA | Quantification of host DNA by qPCR |
